# Applications of machine learning to solve genetics problems

**DOI:** 10.1101/2020.10.27.354092

**Authors:** Kehinde Sowunmi, Victor Nnanna Nweze, Soyebo Titilayo Abiola, Okosesi Ebunoluwa Ajibike, Adesiyan Ayobami Lawal, Olademeji Kafayat Adejoke, Ajibola Oluwaseun Adekunle, Yusuf Olayinka Ogunlana, Agboola Olamilekan Wasiu, Gurpreet Kaur, Atoromola Hameed, Oladipupo Temitope Abidemi

## Abstract

The development of precise DNA editing nucleases that induce double-strand breaks (DSBs) - including zinc finger nucleases, TALENs, and CRISPR/Cas systems - has revolutionized gene editing and genome engineering. Endogenous DNA DSB repair mechanisms are often leveraged to enhance editing efficiency and precision. While the non-homologous end joining (NHEJ) and homologous recombination (HR) DNA DSB repair pathways have already been the topic of an excellent deal of investigation, an alternate pathway, microhomology-mediated end joining (MMEJ), remains relatively unexplored. However, the MMEJ pathway’s ability to supply reproducible and efficient deletions within the course of repair makes it a perfect pathway to be used in gene knockouts. (Microhomology Evoked Deletion Judication EluciDation) may be a random forest machine learning-based method for predicting the extent to which the location of a targeted DNA DSB are going to be repaired using the MMEJ repair pathway. On an independent test set of 24 HeLa cell DSB sites, MEDJED achieved a Pearson coefficient of correlation (PCC) of 81.36%, Mean Absolute Error (MAE) of 10.96%, and Root Mean Square Error (RMSE) of13.09%. This performance demonstrates MEDJED’s value as a tool for researchers who wish to leverage MMEJ to supply efficient and precise gene knock outs.

## Introduction

One of the foremost important applications of gene editing technology is that the creation of gene knockouts for understanding gene function; as of April 01, 2019, quite 11,800 articles indexed in PubMed Central contain the word “knockout” in their title, and therefore the overwhelming majority describe or investigate phenotypes related to gene knockouts [PubMed Central search]. additionally to basic research on gene function, gene knockouts are of critical importance in drug discovery and clinical applications. The creation of gene knockouts that recapitulate human disease phenotypes in model organisms, like mice or zebrafish, allows for rapid and large-scale small molecule and chemical library screening to spot potential therapeutic agents [Parng et al. 2002, Dinda & Baraban 2015, Zuberi & Lutz 2016, Kithcart MacRae 2017, Gehrig et al. 2018]. Gene knockouts have promising clinical applications in gene therapies to directly treat diseases. for instance, tri-nucleotide repeat extension disorders (e.g., Huntington’s chorea [Shin et al. 2016]) could potentially be treated by knocking out the defective protein or excising repeated regions [Cox et al. 2015]. Several ongoing clinical studies are underway to look at the efficacy of knocking out the CCR5 gene (which produces the receptor that macrophage-tropic HIV-1 uses to infect CD4+ T cells) in patient T cells [Case Western Reserve, City of Hope Medical].The ability to induce a double-strand break (DSB) during a gene using targeted nucleases like CRISPR/Cas9 systems [Mali et al. 2013], transcription activator-like effector nucleases (TALENs) [Boch 2011], or zinc finger nucleases (ZFNs) [Kim et al. 1996], [reviewed in Chandrasegaran & Carroll 2016] then leverage endogenous DNA repair pathways to knockout the gene through the method of repair, is very desirable.

## Materials and Methods

### Dataset

Bae et al. kindly provided a dataset of 92 DNA sequences targeted by CRISPR/Cas9 nucleases for DSB [Bae et al. 2014]. All DSB targets were generated in HeLa cells. the info included the gRNA wont to target the DNA sequence, and sequencing read data for every target. The read data had already undergone internal control and was de-duplicated, to get an inventory of the unique amplicon sequences for every target along side a read count for the amplicon.

### Dataset processing

We wrote an R script called Method for Assorting Amplicons Tidily, or MAAT, to sort the recovered amplicons for every target site and characterize features including the amount of wildtype reads, single base insertions and deletions, deletions matching an MMEJ deletion pattern, deletions not matching an MMEJ deletion pattern, and total deletions. Because i used to be not given an inventory of wildtype sequences until much later, we designed MAAT to also plan to identify the canonical wildtype sequence and any SNP variants. It does this by identifying the longest sequence(s) with the most important read count, then identifying single nucleotide differences between the ‘canonical’ sequence and other amplicons of identical length. If wildtype sequences are known, MAAT can use these sequences rather than attempting to spot the wildtype sequence from the read data. During this process, three genes (RP3, CCR5, FLT3) were discarded thanks to low read counts and/or problems with the proto-spacer motif. Unfortunately, MAAT doesn’t (yet) identify cases during which an imperfect microhomology arm match was used. it’s possible for extended microhomology arms to tolerate 1-2 bp mismatches between the arms [Allen] – e.g., ‘GCCACCG’ might pair with ‘CGGGGGC’, albeit the middle nucleotide is mismatched. Although this doesn’t appear to occur frequently and requires long microhomology arms when it does occur [Allen Arbab et al. 2018], it’s possible that MAAT missed a number of these instances within the Bae et al. dataset.

### Training and Validation Datasets

We randomly divided the data into a training set of 75% of the data (66 gene targets) and a validation set of the remaining 23 targets. The IDs of the genes in the training and testing sets is provided in **Table 1**.

**Table-1.**
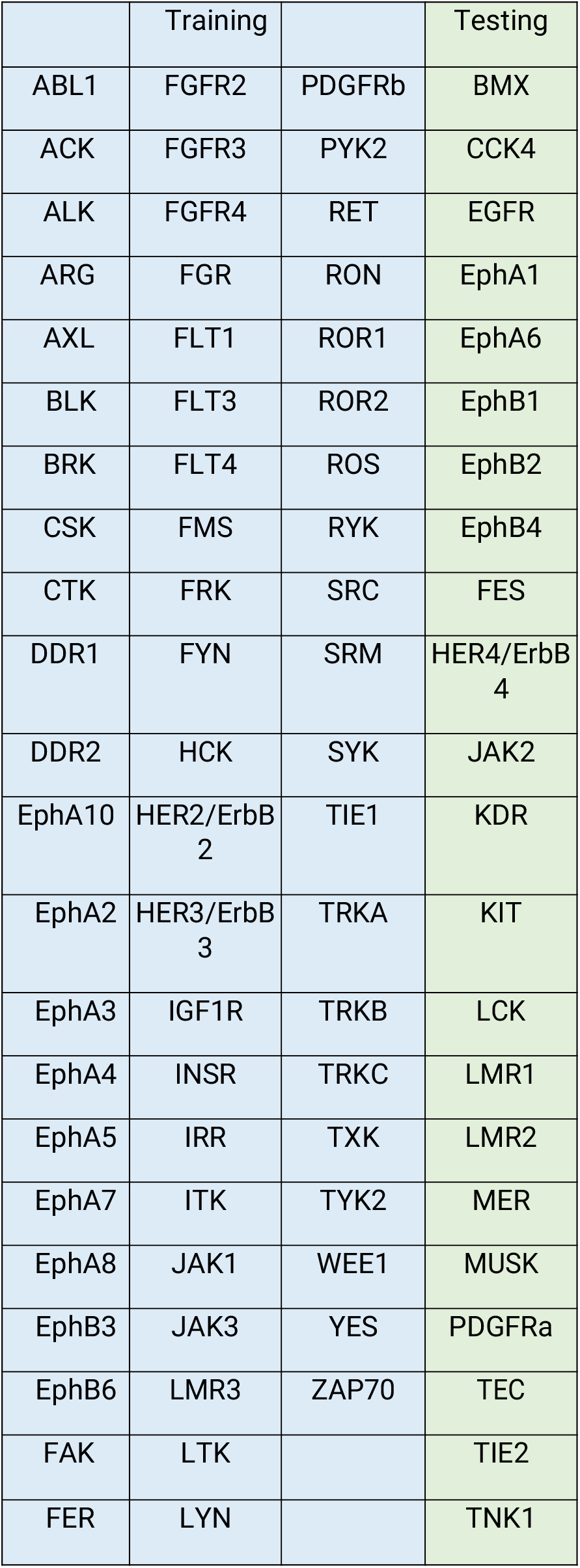
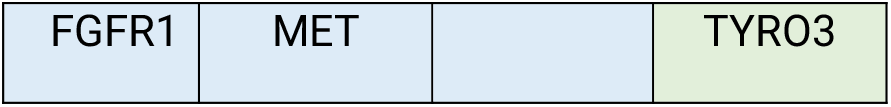
Training and Test set gene IDs

### Dataset Effects

To determine the effect of the dataset on the model, we also randomly sampled 75% of the info, trained a classifier on these data, then tested performance on the remaining 25%. We did this 5000 times and examined the variance in performance metrics.

### Features

Based on the study performed by Bae et al. [Bae et al. 2014] and various features our group determined could be predictive, we generated an initial input vector of 29 features. These features included several “aggregate” features – i.e., features that aggregated information from all potential MMEJ repair patterns for a specific target. These aggregate features included the calculated minimum, maximum, mean, median, and variance of: 1) the amount of nucleotides deleted (deletion length), 2) the length of themicrohomology arm (microhomology arm length), 3) the GC content (number of guanines and cytosines divided by the entire number of nucleotides) of the microhomology arm (microhomology GC content), 4) the amount of nucleotides between the 3’ end of the 5’ microhomology arm and therefore the DSB site (distance to chop site), and 5) the Bae et al. pattern score, for a complete of 25 features. We also assessed a further four features: 1) the GC content of the wildtype DNA sequence surrounding the DSB site, 2) the amount of deletion patterns producing an out-of-frame deletion divided by the entire number of deletion patterns, the Bae microhomology score, and 4) the Bae out-of-frame score.

### Model development

We trained several classifiers using various classification algorithms including Naïve Bayes, logistic regression, generalized linear model, and random forest (not shown). Of these, the random forest classifier worked best. We used the R randomForest package to create the model and perform feature selection.

### Feature selection

Feature importance was assessed using the R randomForest package. thanks to the tiny size of the dataset relative to the dimension of features assessed, we performed feature selection to enhance the model. The randomForest package assesses two measures of importance for regression models: 1) Mean increase in squared error (%IncMSE), which assesses the effect of randomly permuting a feature’s value on the error of the out-of-bag instances, and 2) the decrease in node impurity (IncNodePurity, assessed using the residual sum of squares) caused by splitting the info employing a particular feature (averaged over all the trees within the random forest), which assesses how “close” employing a particular feature gets the regression model to the observed value [Liaw & Wiener 2002]. If randomly permuting a feature produces an outsized increase in MSE, or if it produces an outsized decrease in node impurity, then the feature is vital to the model’s accuracy. Feature selection was performed by assessing feature importance, discarding features with negative %IncMSE (that is, randomly permuting these features improved model performance) and IncNodePurity of 0. a replacement model was created from the remaining features, and this process repeated iteratively until no features had negative %IncMSE and IncNodePurity of 0.

## Results and Discussion

The best-performing model, hereafter mentioned as “MEDJED”, uses 6 features, a forest of 5000 trees, random feature pool of two, and imposes a maximum node constraint of nine nodes per tree. On the training set, it achieved a Pearson coefficient of correlation (PCC) of 81.36%, mean absolute error (MAE) of 10.96%, and root mean squared error (RMSE) of 13.09%. On the test set (which doesn’t overlap with the training set), MEDJED achieved PCC of 85.20%, MAE of 10.26%, and RMSE of 12.02% (see Fig. 4-1). The “zeroR” prediction method, which “learns” the mean of the training data and predicts that value for the test data, achieved an MAE of 18.98%, RMSE of 20.99%, and can’t produce a PCC thanks to the shortage of a typical deviation.

Interestingly, the quality deviation of the pattern score is additionally a crucial feature. Ata et al. formulated a “competition hypothesis” regarding Bae pattern scores and outcomes observed in gene editing experiments - i.e., if there are several microhomologies of comparable “strength”, represented by similar pattern scores, then MMEJ appears to possess trouble “deciding” which microhomology to use, and an alternate pathway might then repair the DSB instead [Ata et al. 2018]. If this is often indeed the case, if the quality deviation of the pattern scores at a specific site is smaller (representing similar pattern scores), then we might expect to ascertain a decrease within the MMEJ at that site.I plotted the quality deviation of the pattern score for every target site against the observed proportion of MMEJ-based deletions, and located a PCC of 56.47% (see Fig. 2), which shows that increasing variance of the pattern scores at a specific deletion site correlates moderately well with its observed MMEJ deletion outcomes, thus providing support for the Ata et al. competition hypothesis.The correlation between the quality deviation of the microhomology arm lengths and repair outcomes (PCC 41.53%) lends further support to the competition hypothesis: a extended microhomology arm results in a bigger pattern score, then if there’s large variation in microhomology arm lengths, this suggests large variation in microhomology arm strengths, and thus less competition for one predominant MMEJ outcome, remaining features are the mean and maximum microhomology arm lengths, which correlate moderately well with the observed MMEJ repair outcomes (PCC 40.00% and 33.04%, respectively.) The importance of the mean and maximum microhomology arm lengths is probably going thanks to MMEJ favoring long microhomologies for base pairing [Allen & Crepaldi et al. 2018, Shen & Arbab et al. 2018].

**Figure-1.**
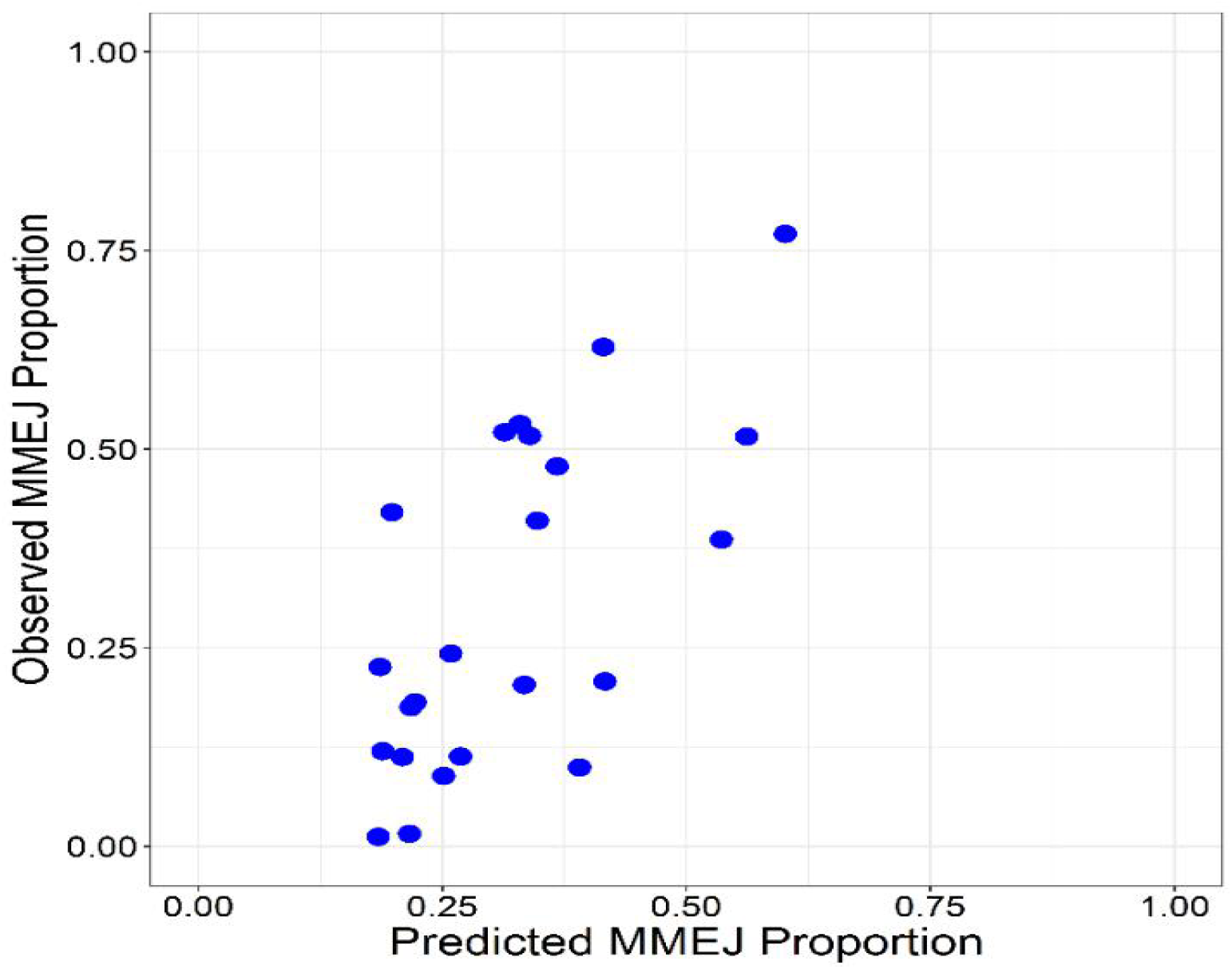
MEDJED performance. On the independent test set, MEDJED achieves a Pearson Correlation Coefficient (PCC) of 81.36%, Mean Absolute Error (MAE) of 10.96%, and Root Mean Square Error (RMSE) of 13.09%. The MEDJED-predicted MMEJ repair proportion (x-axis) is graphed against the observed MMEJ repair proportion (y-axis).

**Figure 2.**
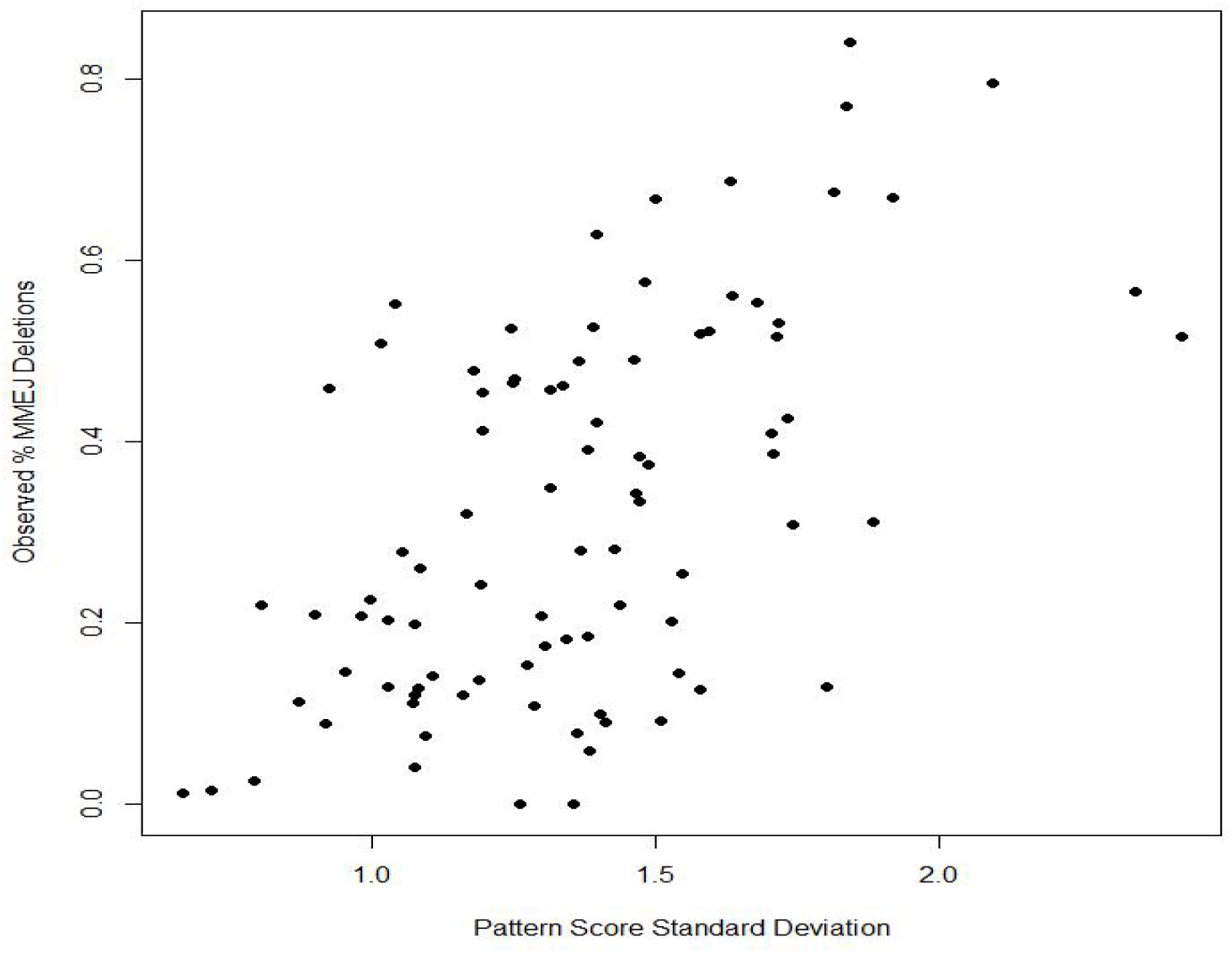
Scatterplot of the standard deviation of the Bae et al. [Bae et al. 2014] pattern score plotted against the observed proportion of MMEJ deletions for 89 HeLa cell targets. Pearson correlation coefficient of 56.47%.

### Dataset Effects

To determine if the model’s performance was a results of a fortunate split between examples within the training and testing set, I randomly divided the HeLa dataset into training and test sets 5000 times then trained and tested random forest classifiers with MEDJED’s parameters (see Fig. 3). The mean absolute error (MAE) for these 5000 trees ranged from 8.73% to 19.80%, with median 13.97% and mean of 14.02% compared to the ultimate MEDJED model’s MAE of 10.26%. the basis mean square error (RMSE) of the 5000 trees ranged from 10.57% to 22.43%, with median 16.54% and mean 16.56%, compared to MEDJED’s RMSE of 12.02%. The Pearson Correlation Coefficients of the 5000 trees ranged from just 11.20%to 91.03%, compared to MEDJED’s PCC of 85.20%.This demonstrates that while the ultimate MEDJED model may enjoy some effectsof the random split of the training and test data, the model isn’t a statistical outlier.

**Figure 3.**
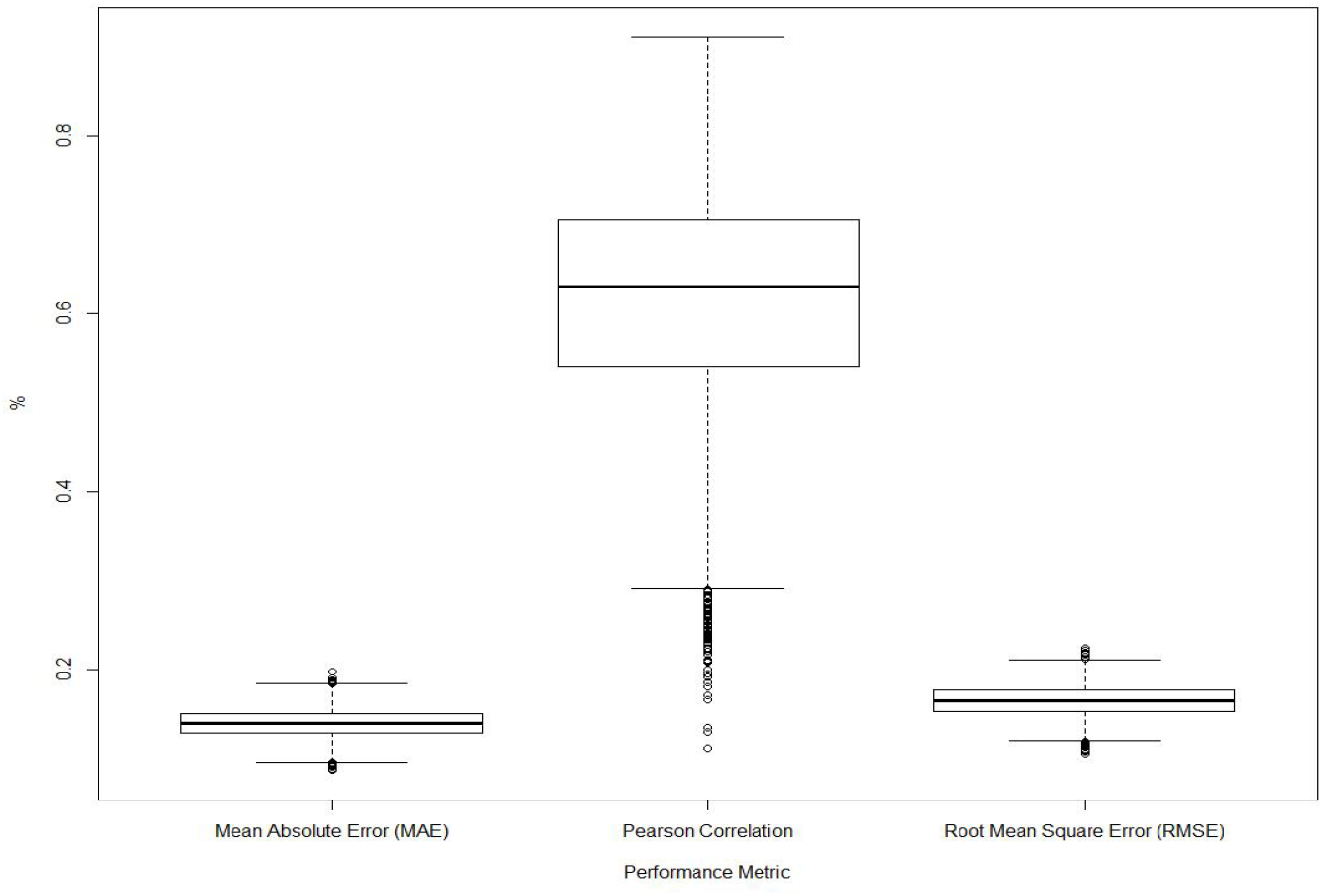
Boxplot of the mean absolute error (MAE), Pearson Correlation Coefficient (PCC), and root mean squared error (RMSE) on test sets. The original dataset was randomly split into training (75%) and testing sets (25%) 5000 times, generating 5000 classifiers with the same parameters as the final MEDJED model; each classifier was then assessed on its respective test dataset. The median MAE was 13.97% with standard deviation of 1.6%. The median PCC was 63.06%, with standard deviation 12.39%. The median RMSE was 16.54% with standard deviation of 1.7%.

## Conclusion

MEDJED may be a machine learning regression model for predicting the extent to which a DSB site will undergo microhomology-mediated end joining. MEDJED is capable of predicting the extent of MMEJ-repair utilization in sequence deletion outcomes, and is out there online at www.genesculpt.org/medjed/, and may be downloaded at https://github.com/Dobbs-Lab/medjed/. it’s also available as a Docker image at https://hub.docker.com/r/cmmann/medjed.

In November 2018 (several months after the event of MEDJED), a gaggle from the Wellcome Sanger Institute [Allen & Crepaldi et al. 2018] and a gaggle including scientists from the Broad Institute, the Massachusetts Institute of Technology, and Harvard school of medicine [Shen & Arbab et al. 2018] both published methods, called FORECasT and inDelphi, respectively, for predicting the sequence of repair outcomes at DSB sites. The FORECasT method was demonstrated to realize near the theoretical limit of prediction accuracy (i.e., the extent of reproducibility between gene editing experiments) [Allen & Crepaldi et al. 2018]. FORECasT was trained using ~40,000 DSB sites constructed for the study [Allen and Crepaldi et al. 2018]; inDelphi was trained with ~4000 DSBs similarly generated to be used within the study [Shen & Arbab et al. 2018]. The existence of those tools makes the further development of MEDJED to predict the sequence of repair outcomes unnecessary and is of much more interest, however, is that the potential MEDJED offers for understanding the biological mechanisms of DNA DSB repair. FORECasT requires enumerating all possible deletions and short insertions.

## Acknowledgements

We would like to acknowledge and thank the Central Lab group, especiallyl for their assistance in bug-testing and hosting early versions of MEDJED, as well as valuable discussions. We would also like to thank the Suliat Shoga and Ibadu Rahman labs for valuable discussions and bug testing.

## References

Allen, F., Crepaldi, L., Alsinet, C., Strong, A.J., Kleshchevnikov, V., De Angeli, P., Palenikova, P., Khodak, A., Kiselev, V., Kosicki, M. et al.(2019) Predicting the mutations generated by repair of Cas9-induced double-strand breaks. Nature Biotechnology, 37, 64–72. http://dx.doi.org/10.1038/nbt.4317

, H., Ekstrom, T.L., Martinez-Galvez, G., Mann, C.M., Dvornikov, A.V., Schaefbauer, K.J., Ma, A.C., Dobbs, D., Clark, K.J. and Ekker, S.C. (2018) Robust activation of microhomology-mediated end joining for precision gene editing applications. PLoS Genetics, 14, e1007652. http://dx.doi.org/10.1371/journal.pgen.1007652

Bae, S., Kweon, J., Kim, H.S. and Kim, J.S. (2014) Microhomology-based choice of Cas9 nuclease target sites. Nature Methods, 11, 705–706. http://dx.doi.org/10.1038/nmeth.3015

Bétermier, M., Bertrand, P. and Lopez, B.S. (2014) Is non-homologous end-joining really an inherently error-prone process? PLoS Genetics, 10, e1004086.http://dx.doi.org/10.1371/journal.pgen.1004086

Boch, J. (2011) TALEs of genome targeting. Nature Biotechnology, 29, 135–136. http://dx.doi.org/10.1038/nbt.1767

Case Western Reserve, U., University of California, S.F. and University of, C.(2024). CCR5-modified CD4+ T Cells for HIV Infection. NCT03666871.

Chandrasegaran, S. and Carroll, D. (2016) Origins of Programmable Nucleases for Genome Engineering. Journal of Molecular Biology, 428, 963–989. http://dx.doi.org/10.1016/j.jmb.2015.10.014

City of Hope Medical, C., Sangamo, T. and California Institute for Regenerative, M. (2019). Safety Study of Zinc Finger Nuclease CCR5-modified Hematopoietic Stem/Progenitor Cells in HIV-1 Infected Patients. NCT02500849.

Cox, D.B., Platt, R.J. and Zhang, F. (2015) Therapeutic genome editing: prospects and challenges. Nature Medicine, 21, 121–131.http://dx.doi.org/10.1038/nm.3793

Dinday, M.T. and Baraban, S.C. (2015) Large-Scale Phenotype-Based Antiepileptic Drug Screening in a Zebrafish Model of Dravet Syndrome. eNeuro, 2, epublish. http://dx.doi.org/10.1523/eneuro.0068-15.2015

Gehrig, J., Pandey, G. and Westhoff, J.H. (2018) Zebrafish as a Model for Drug Screening in Genetic Kidney Diseases. Frontiers in Pediatrics, 6, 183. http://dx.doi.org/10.3389/fped.2018.00183

Thomas, K.R. and Capecchi, M.R. (1987) Site-directed mutagenesis by gene targeting in mouse embryo-derived stem cells. Cell, 51, 503–512.http://www.ncbi.nlm.nih.gov/pubmed/2822260

van Overbeek, M., Capurso, D., Carter, M.M., Thompson, M.S., Frias, E., Russ, C., Reece-Hoyes, J.S., Nye, C., Gradia, S., Vidal, B. et al.(2016) DNA Repair Profiling Reveals Nonrandom Outcomes at Cas9-Mediated Breaks. Molecular Cell, 63, 633–646. http://dx.doi.org/10.1016/j.molcel.2016.06.037

Vasquez, K.M., Marburger, K., Intody, Z. and Wilson, J.H. (2001) Manipulating the mammalian genome by homologous recombination. Proceedings of the National Academy of Sciences of the United States of America, 98, 8403–8410.http://dx.doi.org/10.1073/pnas.111009698

Zuberi, A. and Lutz, C. (2016) Mouse Models for Drug Discovery. Can New Tools and Technology Improve Translational Power? ILAR Journal, 57, 178–185. http://dx.doi.org/10.1093/ilar/ilw021

